# MOBFinder: a tool for MOB typing for plasmid metagenomic fragments based on language model

**DOI:** 10.1101/2023.12.06.570414

**Authors:** Tao Feng, Shufang Wu, Hongwei Zhou, Zhencheng Fang

## Abstract

**Background:** MOB typing is a classification scheme that classifies plasmid genomes based on their relaxase gene. The host range of plasmids of different MOB categories are diverse and MOB typing is crucial for investigating the mobilization of plasmid, especially the transmission of resistance genes and virulence factors. However, MOB typing of plasmid metagenomic data is challenging due to the highly fragmented characteristic of metagenomic contigs.

**Results:** We developed MOBFinder, an 11-class classifier to classify the plasmid fragments into 10 MOB categories and a non-mobilizable category. We first performed the MOB typing for classifying complete plasmid genomes using the relaxes information, and constructed the artificial benchmark plasmid metagenomic fragments from these complete plasmid genomes whose MOB types are well annotated. Based on natural language models, we used the word vector to characterize the plasmid fragments. Several random forest classification models were trained and integrated for predicting plasmid fragments with different lengths. Evaluating the tool over the benchmark dataset, MOBFinder demonstrates higher performance compared to the existing tool, with an overall *accuracy* of approximately 59% higher than the MOB-suite. Moreover, the *balanced accuracy, harmonic mean* and *F1-score* could reach 99% in some MOB types. In an application focused on a T2D cohort, MOBFinder offered insights suggesting that the MOBF type might accelerate the antibiotic resistance transmission in patients suffering from T2D.

**Conclusions:** To the best of our knowledge, MOBFinder is the first tool for MOB tying for plasmid metagenomic fragments. MOBFinder is freely available at https://github.com/FengTaoSMU/MOBFinder.

## 1. Introduction

Plasmids are usually small, double-stranded, and circular DNA molecules found within bacterial cells [1]. Existing separately from the bacterial chromosome, plasmids have the ability to replicate independently and can transition between bacteria through conjugation [2]. Bacteria, specifically pathogenic strains, can acquire antibiotic resistant genes or virulence factors via plasmid-mediated horizontal gene transfer, thereby equipping them to adapt to various environments [3].

Plasmid classification is important for investigating multiple properties of plasmids, such as host range, replication and mobilization mechanism [4]. Many classification schemes have been developed according to the distinct characteristics of plasmids, like taxonomic classification, replicon (Rep) typing, incompatibility (Inc) typing, mate-pair formation (MPF) typing and mobilization (MOB) typing. Taxonomic classification refers to classify plasmids based on their host bacteria [5]. Rep typing is accomplished by genes controlling plasmid replication, known as replicons [4, 6]. As plasmids with similar replication or partition system are incompatible within the same cell, Inc typing is a method of categorizing plasmids based on their compatibility [6]. MPF typing is based on the discovery of genes encoding the MPF system, which consists of proteins that can mediate contact and DNA exchange between donor and recipient cells during conjugation [4, 7]. Compared to these methods, MOB typing, another classification scheme, classifies plasmids based on the relaxase gene, which is present in all transmissible plasmids [8-10]. Plasmids with different MOB types, classified according to their relaxase types, possess dinstinct transmission mechanisms that determine their taxonomic host range [4, 11]. These variations among different MOB types are critical in researching the spread of virulence traits, the emergence of antibiotic resistance, and the adaption and evolution of bacteria. Moreover, MOB typing has demonstrated its effectiveness in identifying novel mobilizable plasmids that were previously unassigned to any Rep or Inc types, and investigating the mobilization characteristics of plasmids that have similar mobilization systems [12, 13].

Recently, many experimental and computational schemes have been devised for plasmid typing, as well as to explore the diversity and functionality of plasmids (Table 1). PlasTax-PCR (PLASmid TAXonomic PCR) [14], PBRT (PCR-Based Replicon Typing) [15], and DPMT (Degenerate Primer MOB Typing) [12] are multiplex PCR methods devised to identify plasmids with analogous replication or mobilization systems. PlasTrans, designed based on deep learning, was available to identify mobilizable metagenomic plasmid fragments [16]. Web servers like PlasmidFinder [6], pMLST, and oriTfinder [17] were established utilizing collected maker gene databases and alignment-based methods, to facilitate Rep typing, Inc typing, or MOB typing. COPLA [5], developed based on average nucleotide identity, is a tool designed to perform taxonomic classifications of complete plasmid genomes with an overall accuracy of 41%. Unlike these methods, MOB-suite [18, 19] was designed to perform plasmid typing for plasmid assemblies. MOB-suite was developed to achieve plasmid typing of plasmids assemblies. MOB-suite using mash to cluster plasmid assemblies into clusters, and then using collected marker gene databases to annotate them with an *e-value* of 1e-5, a *query coverage* of 80% and an *identity* of 80%.

**Table 1.** Experimental and computational schemes developed for plasmid classification.

| Technology category | Method | Classification scheme | Material | Description |
| --- | --- | --- | --- | --- |
| Experimental | DPMT [12] | MOB typing | Plasmid DNA collections from clinical isolates | Using degenerate primers to hybridize relaxase-coding gene to identify and classify plasmids isolated from clinical isolates |
|  | PlasTax-PCR [14] | Taxonomic typing | Plasmid DNA collections from clinical isolates | Utilized PCR primers that target conserved segments of the relaxase gene of plasmid taxonomic units (PTUs) to identify specific PTUs of transmissible plasmids |
|  | PBRT [15] | Rep typing or Inc typing | Plasmid DNA collections from clinical isolates | Used multiplex PCR to amplify DNA fragments of replicon and detect known replicon types of plasmids |
| Computational | MOB-suite [18, 19] | MOB typing, MPF typing and Rep typing | Complete plasmid genomes or plasmid assembly clusters (Linux) | Utilized collected relaxase, oriT, replicon and T4SS sequences to construct database, then performing classification for plasmids assembly clusters with BLAST |
|  | PlasTans [16] | transmissible plasmid identification | plasmid assembly contigs (Linux) | Using the convolutional neural network of the deep learning technique to classify plasmid DNA fragment |
|  | PlasmidFinder [6] | Rep typing or Inc typing | Raw reads or complete plasmid genomes or plasmid assembly contigs (web server) | Utilized collected replicon sequences and BLASTn to perform Rep typing and Inc typing |
|  | pMLST [6] | Rep typing or Inc typing | Raw reads or Complete plasmid genomes or plasmid assembly contigs (web server) | Used collected plasmid multilocus sequence typing (pMLST) allele sequences, known sequence type profiles and BLAST to perform Rep typing and Inc typing |
|  | oriTfinder [17] | MOB typing, MPF typing | Complete plasmid genomes (web server) | Utilized collected oriT, relaxase, T4CP and T4SS sequences to annotate plasmids with BLAST |
|  | COPLA [5] | Taxonomic typing | Complete plasmid genomes or plasmid assembly sets (Linux) | Using average nucleotide identity (ANI) metrics and hierarchical stochastic block modeling (HSBM) to create plasmid taxonomic units (PTUs) and predict the taxonomic host |

Metagenomic sequencing makes it possible to obtain all plasmid DNA from microbial communities at once, and a number of computational tools for identify plasmid fragments from metagenomic data have been developed, such as PlasFlow [20], PlasmidSeeker [21], PlasClass [22], PPR-Meta [23] and PlasForest [24]. As the DNA fragments of plasmids and bacteria are intermingled in metagenomic data [25, 26], recognizing the host and transmission range of plasmids can be challenging. To explore the host range and transmission mechanism of plasmids with different mobilizable systems using metagenomic sequencing, it is crucial to achieve the MOB annotation of metagenomic plasmid fragments. However, obstacles arise due to the incompleteness of plasmid assembly fragments from metagenomic data and the absence of essential genes for annotation, thereby making it difficult to accurately annotate the MOB class of plasmid fragments. Given that plasmids with the same MOB type share similarities in their transmission mechanisms and host ranges, the genomic signatures, such as the GC content and codon usage of each MOB type tend to be alike, not only relaxase [4, 27]. Neural networks have demonstrated powerful performance in the classification and identification of biological sequences [28, 29]. Furthermore, language models [30, 31] derived from these neural networks have also showcased their impressive ability to characterize sequence features [32, 33]. To characterize the features of plasmids within the same MOB type, we employed language models to perform the MOB annotation. In addition to the relaxase-coding gene, language models exhibit the ability to capture more biological features and associations within comparable mobilization systems, making it possible to perform MOB annotation for metagenomic plasmid assemblies.

To address the challenge of MOB typing for metagenomic plasmid fragments, we presented MOBFinder, a tool designed for annotating MOB types in plasmid metagenomic fragments. MOBFinder can process single or multiple plasmid DNA sequences, and provide predicted MOB types for input each fragment, including MOBB, MOBC, MOBF, MOBH, MOBL, MOBM, MOBP, MOBQ, MOBT, MOBV and non-mob. Moreover, MOBFinder also provides the option to annotate plasmid bins from metagenomics data. The development of MOBFinder involved the following steps: (1) Benchmark dataset construction: Plasmid complete genomes obtained from National Center for Biotechnology Information (NCBI) were classified into different MOB types based on relaxase databases. To simulate plasmid fragments in metagenomic data, artificial benchmark datasets with varying lengths were generated. (2) Word embeddings. Numerical word vectors were generated using skip-gram to characterize the sequence features in different MOB categories. (3) Classification model ensemble and optimization. Several classification models, specifically designed for different lengths, were trained and then integrated to enhance the overall performance of MOBFinder. The evaluation of the test dataset demonstrated that MOBFinder is a powerful tool for MOB typing of plasmid fragments and bins. MOBFinder’s application in a T2D cohort revealed a potential correlation between the MOBF type and the spread of antibiotic resistance genes among T2D patients. This highlights the potential significance of MOBFinder in understanding and addressing antibiotic resistance in specific patient populations.

## 2. Materials and methods

### 2.1. The workflow of MOBFinder

To annotate the MOB type of plasmid fragments in metagenomics, we designed MOBFinder (Figure 1). As MOB-suite [18, 19] didn’t offer a quantitative likelihood score for the outcomes and some plasmids would be classified into multiple MOB types (Figure S1), we constructed a benchmark dataset using a high-resolution MOB typing strategy to for categorizing complete plasmid genomes (Figure 1A). Then, based on a language model and random forest, we designed an algorithm to perform MOB typing for plasmid metagenomic fragments (Figure 1B).

**Figure 1.**
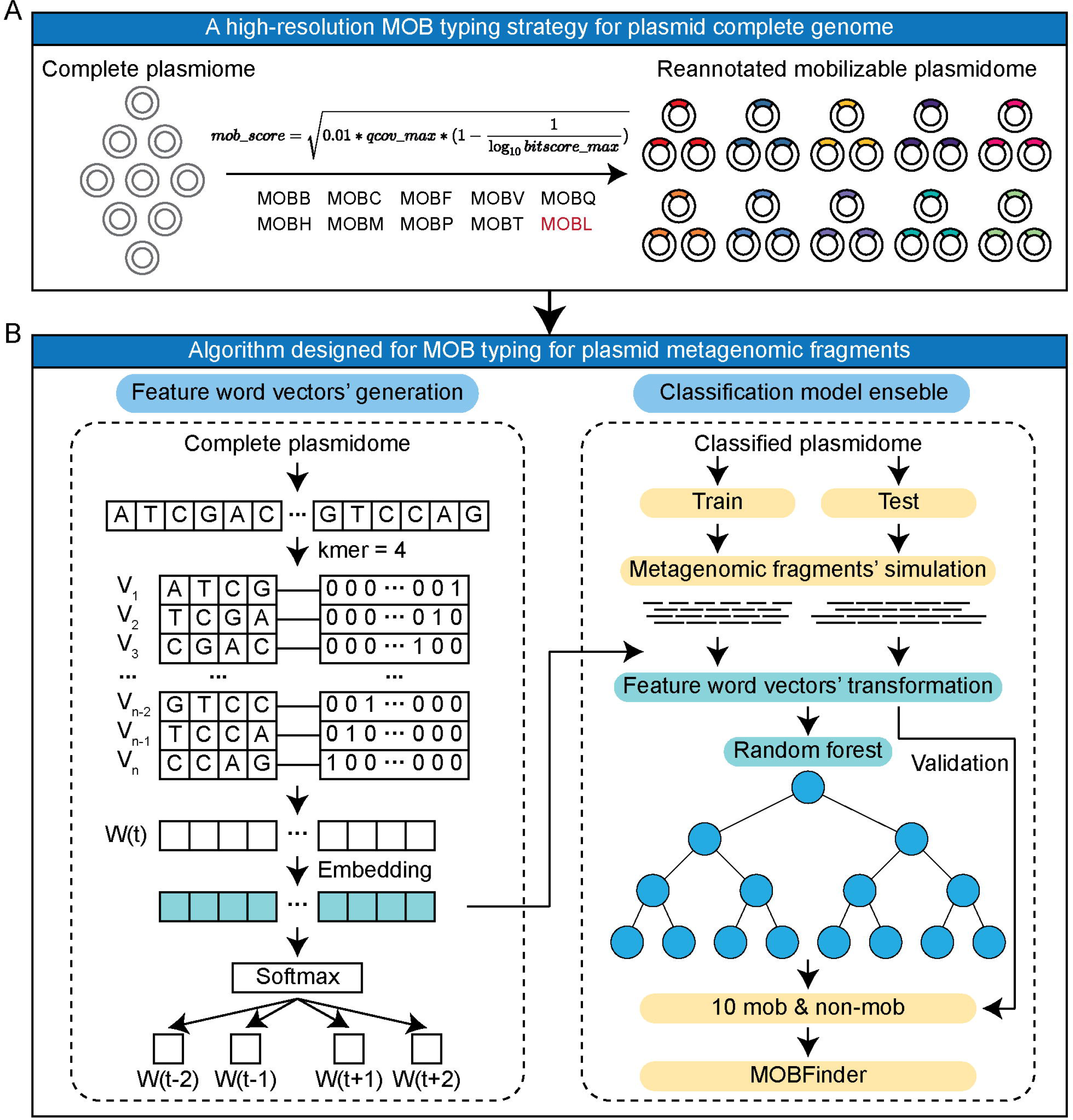
The workflow of MOBFinder. (A) The benchmark dataset was created using a relaxase-based method, employing a high-resolution strategy to classify different MOB types of complete plasmid genomes. This classification relied on a combination of defined *mob_score* and *evalue*. (B) The algorithm designed for MOBFinder consisted of two main modules. a. Feature word vectors were generated to characterize the plasmids of MOB types. b. Several random forest classification models were integrated to predict plasmid metagenomic fragments with different lengths.

### 2.2. MOB typing for complete plasmid genomes

Ten validated MOB relaxase protein families were collected, including MOBB, MOBC, MOBF, MOBH, MOBL, MOBM, MOBP, MOBQ, MOBT and MOBV [7-10, 34-37]. For each MOB category, blastp [38] was used to search homologous protein sequences against NCBI non-redundant protein sequence database, with an *evalue* threshold of 1e-10, a *query coverage* threshold of 70% and an *identity* threshold of 70%. After the expansion of protein sequences, local relaxase databases were built using the ‘makeblastdb’ command for MOB typing of plasmid genomes.

Plasmid genomes were retrieved from the NCBI nucleotide database using the keywords ‘complete’ and ‘plasmid’, and incomplete plasmid fragments were removed manually for further analysis. The accession list of these plasmids is provided in Supplementary Table 1. For each plasmid genome, coding sequences were extracted from the genebank file, and blastp [38] was employed to search for the best alignment of local relaxase databases. Here, we defined the *mob_score* (a) to measure the likelihood of homology:

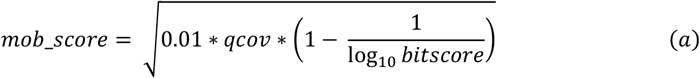

where *qcov* and *bitscore* represent the *query coverage* and *bitscore* corresponding to the best match, respectively. To identify plasmid genomes encoding known relaxase families, we set a *mob_score* threshold of 0.5, which was established in conjunction with a minimum *query coverage* of 50 and a minimum *bitscore* of 100. To further enhance the reliability of our classification, we introduced an evalue cutoff, conservatively set at 1e-10, to facilitate the plasmid genome classification. In instances where plasmid genomes yielded no blast results or exhibited an *evalue* exceeding 0.01, we categorized them as non-mob.

### 2.3. Word embeddings using a language model

To characterize the features and patterns within each MOB category and use numerical word vectors to represent them, we utilized a skip-gram language model [30, 31] to learn from plasmid genomes. Using a fixed-size sliding window, the skip-gram algorithm calculated the likelihood between segmented words and outputted a probability distribution over the context words. The training steps were as follows:

(1) Word generation. Because the plasmid DNA sequences are composed of four nucleotide bases: ‘A’, ‘T’, ‘C’, and ‘G’, we used a 4-mer sliding window to generate overlapping input words. For example, ‘ATCGCTGA’ would be segmented into ‘ATCG’, ‘TCGC’, ‘CGCT’, ‘GCTG’ and ‘CTGA’. In this step, 256 unique words will be generated.
(2) Word encoding. One-hot encoding was utilized to represent input words, and these words would be transformed to a 256-dimensional vector consisting of 255 zeros and a unique 1.
(3) Skip-gram model. A two-layer neural network was used to construct the skip-gram model. The input is one-hot encoding vectors, and the output is a probability distribution over the input words. Layer 1 is a hidden layer to convert the input one-hot encoding into a 100-dimensional word vector representation. Layer 2 is used to compute and maximize the probability of the correct context words using the softmax function, and the size of context words was set to 20.
(4) Model training. For each input plasmid genome, we used an optimization algorithm to minimize the loss function. Then we used backpropagation to update the neural network parameters (word vectors) until convergence.
(5) Word vector extraction. After the training process, the word vectors in the hidden layer would be extracted to characterize the plasmid fragments in metagenomics.

### 2.4. Benchmark datasets construction

Because there is no real metagenomic data to serve as a benchmark [16, 23], we artificially generated simulated datasets with the following steps:

(1) For classified plasmid genomes in each MOB category, we randomly split them with a proportion of 70% and 30% to construct the training and test datasets.
(2) Training dataset. To predict plasmid fragments with different lengths, we generated contigs with different length ranges: 100-400 bp, 401-800 bp, 801-1,200 bp and 1,201-1,600 bp. For each MOB class in four length ranges, we randomly generated 90,000 artificial contigs. Plasmid fragments longer than 1600 bp would be segmented into shorter contigs and predicted using models designed for the corresponding lengths.
(3) Test dataset. Because the plasmid fragments in real metagenomics were much longer, we generated other four length groups to assess the performance of MOBFinder: Group A with a length range of 801-1,200 bp, Group B with a length range of 1201-1,600 bp, Group C with a length range of 3,000-4,000 bp and Group D with a length range of 5,000-10,000 bp. For each MOB class in four groups, 500 fragments were randomly extracted.

### 2.5. Classification algorithm design

To efficiently handle the training dataset and improve the robustness of MOBFinder, we employed random forest to train four predictive models using the training dataset. The detailed steps are as follows:

(1) Word representations’ calculation. For each contig in the training dataset, we used a 4-mer sliding window to generate overlapping words and transformed them into numerical word vectors using trained word embeddings. To characterize underlying features and patterns of the input contigs, we summed up all the word vectors to compute their average as input of random forest.

(2) Classification models’ training. To improve the performance of MOBFinder, we trained four classification models for different lengths in the training dataset: 100-400 bp, 401-800 bp, 801-1,200 bp and 1,201-1,600 bp, and the number of trees was set to 500 to generate predictive models.

(3) Model ensemble. Four trained models for different lengths were ensembled into MOBFinder to make more accurate predictions. For fragments shorter than 100 bp, we used a model designed for 100-400 bp to predict the MOB type. For fragments longer than 1,600 bp, we segmented them into short contigs according to the length of trained models and make predictions using the corresponding model. For example, a fragment with a length of 4,000 bp would be segmented into two contigs with a length of 1,600 bp and one contig with a length of 800 bp. After predicting with the corresponding models, we aggregated and calculated the average scores for each MOB class, and the MOB type with the highest score will be selected as the predicted result for the input fragment (b). The calculation formula is:

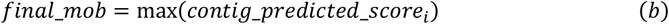

where *contig_predicted_score*_*i*_ represents the predicted score of the input fragment for each MOB type and *final_mob* represents the predicted MOB category with the highest score.

(4) Plasmid bins’ classification. Metagenomic binning is an essential step for the reconstruction of genomes from individual microorganisms. Thus, MOBFinder can perform MOB typing on both plasmid contigs and plasmid bins. If the input is a plasmid bin, MOBFinder predicted the likelihood of each MOB class for fragments within the bin. MOBFinder calculates the score for each MOB type, and selects the MOB type with the highest score as the MOB type for the plasmid bin (c), and the calculation formula is:

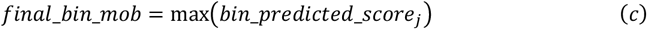

where *bin_predicted_score*_*j*_ represents the predicted scores of the input plasmid bin for each MOB type and *final_bin_mob* represents the predicted MOB category with the highest score.

### 2.6. Performance validation

Test dataset was used to assess the performance of MOBFinder, and compared to MOB-suite [18, 19]. We calculated overall *accuracy, kappa* and *run time* by comparing the predicted classes and true classes. The overall *accuracy* was the proportion of accurate predictions. The *kappa* (d) was calculated to assess the overall consistency between the predictions and true classes, which took into account the possibility of random prediction. *Po* represents observed accuracy [*Po* = (*A*_*11*_ + *A*_*22*_ + … + *Ann*) / *N*], where *A*_*11*_, *A*_*22*_ and *Ann* represent the values on the diagonal of the confusion matrix, and *n* represents the number of MOB categories, while *N* represents the total number of samples. *Pe* represents expected accuracy [*Pe* = (*E*_*11*_ + *E*_*22*_ + … + *Enn*) / *N*^2], where *E*_*11*_, *E*_*22*_ and *Enn* represent the expected values in each cell of the confusion matrix, *n* represents the number of MOB classes, and *N* represents the total number of samples. The *runtime* was recorded using the command ‘time’ in Linux.

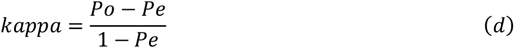

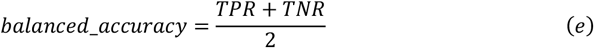

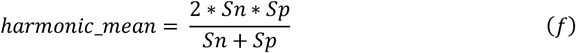

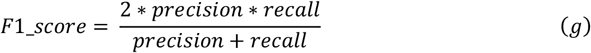

For each MOB category, we also calculated the *balanced accuracy* (e), *harmonic mean* (f) and *F1_score* (g). Considering the class imbalance within the training dataset, *balanced accuracy* was used to measure the average accuracy of each MOB category, where *TPR* is the true positive rate [*TRP* = true positives / (true positives + false negatives)], and *TNR* is the true negative rate [*TNR* = true negatives / (true negatives + false positives)]. *Harmonic mean* could provide an overall evaluation of a model’s performance, where *Sn* and *Sp* represent sensitivity [*Sn* = true positives/ (true positives + false negatives)] and specificity [*Sp* = true negatives / (true negatives + false positives)]. *F1-score* combines *precision* and *recall*, providing a balanced measure of a model’s performance, where *precision* was the correct positive prediction in all positive predictions [*precision* = true positives / (true positives + false positives)] and *recall* was the correct positive predictions in all actual positives [*recall* = true positives / (true positives + false negatives]].

The receiver operating characteristic (ROC) curve was used to visualize the performance of MOBFinder in predicting each MOB category, where the x-axis and y-axis were false positive rate (*FPR*) and true positive rate (*TPR*). The ROC curve that is closer to the left and top indicates a higher *TPR* and lower *FPR*, which means better performance. For each MOB class, the area under the curve (AUC) value was calculated to quantify the performance of MOBFinder. The AUC value between 0.5 and 1 indicates the model performs better than random chance, and a higher AUC value indicates better prediction capability.

### 2.7. Annotation and analysis of T2D metagenomic data

Metagenomic sequencing data (SRA045646) were retrieved from NCBI short read archive (SRA) database to investigate whether the plasmids within different MOB classes were associated with the resistance enrichment in T2D patients, as suggested by a previous study [39, 40]. All metagenomic data were preprocessed using the same protocols. PRINSEQ [41] was used to remove low-quality reads and bowtie2 [42] to remove host reads by aligning them to the human GRCH38 reference genome downloaded from ENSEMBL database. We excluded metagenomic samples that did not pass the quality control. Because the plasmid abundance in metagenomes was much lower than bacteria, we only retained samples with more than 10,000,000 paired-end reads for downstream analysis (Supplementary Table 2).

**Table 2.** The number, average length, and GC content of plasmid genomes for each MOB type.

| Class | Number | Average length | GC (%) |
| --- | --- | --- | --- |
| MOBB | 623 | 10921.77 | 51.27 |
| MOBC | 3218 | 19965.28 | 47.14 |
| MOBF | 21268 | 103802.80 | 52.07 |
| MOBH | 4880 | 151108.10 | 48.37 |
| MOBL | 3446 | 51430.63 | 34.57 |
| MOBM | 1761 | 2684.14 | 27.12 |
| MOBP | 15617 | 32237.88 | 49.70 |
| MOBQ | 9347 | 89357.64 | 56.77 |
| MOBT | 1181 | 11643.24 | 36.92 |
| MOBV | 4405 | 6595.43 | 37.75 |
| Non-mob | 24649 | 37581.85 | 49.84 |

To improve the efficiency and accuracy of assembly, we used MEGAHIT [43] to generate metagenomic fragments. PPR-Meta [23] was utilized to identify and extract plasmid fragments from the assembled fragments, while filtering out bacterial and phage sequences. COCACOLA [44] was employed to cluster plasmid fragments into bins based on their sequence similarity and composition. This could help us to investigate the plasmid fragments from same originate and enable a better annotation and anlysis of their functions.

MOBFinder was applied to annotate the MOB types for each plasmid bin. The average fragments per kilobase per million (FPKM) of each plasmid bin was calculated using bowtie2 to represent its abundance. Using Wilcoxon signed-rank two-sided test, Plasmid bins with a *p-value* less than 0.05 were selected as T2D-related plasmid bins. ABRicate [45] was utilized to annotate the antibiotic resistance genes in each plasmid bin, based on four antibiotic resistance gene databases [46-49]. The Tukey’s Honest Significant Difference test was performed to compare the identified resistant genes for different MOB classes. All statistical analysis were achieved using R.

## 3. Results

### 3.1. MOB typing for plasmid genomes

To construct the benchmark datasets, we obtained 90,395 complete plasmid genomes and categorized them into 11 MOB categories using blast (Table 2). We removed 22,470 plasmid genomes potentially classified into more than one MOB class, leaving 67,925 classified genomes for the training and optimization of MOBFinder (Figure 2A). Our analysis results revealed significant variations among the plasmid genomes of different MOB types, including the number, average length, and GC content. Notably, non-MOB type plasmid genomes held the highest count and longest average length, whereas MOBB and MOBM had the fewest plasmid genomes and shortest average length, respectively. In terms of GC content, MOBL and MOBQ represent the lowest and highest MOB type. Moreover, plasmids of different MOB types exhibited diverse host ranges in genus level (Figure 2B). MOBB was predominantly found in *Hymenobacter, Parabacteroides, Phocaeicola* and *Spirosoma*. Particularly, *Phocaeicola* has been detected in the human gut and possessed the gene for porphyran degradation through horizontal gene transfer [50]. MOBC, MOBF, MOBH and MOBP all existed in *Escherichia* and *Klebsiella*. Furthermore, *Klebsiella* is a multidrug-resistant bacterium that has demonstrated resistance to multiple antibiotics [51]. MOBL, MOBT and MOBV were mainly discovered in *Bacillus* and *Enterococcus*.

**Figure 2.**
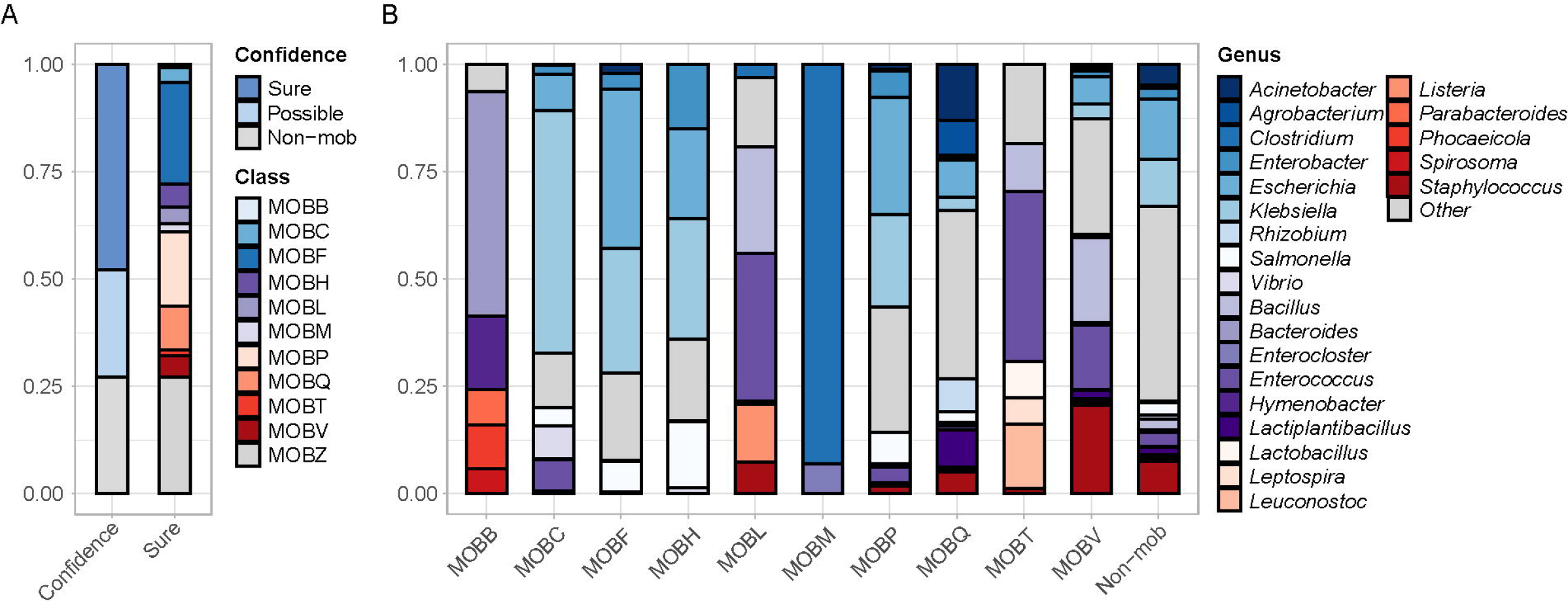
Benchmark dataset construction using a high-resolution strategy. (A) The proportion of classified plasmid genomes. The confidence is ‘sure’ means the classified plasmid genomes had a *mob-score* more than 0.5 and an *evalue* less than 1e-10, while ‘possible’ had not. Plasmid genomes identified as ‘sure’ were used to generate benchmark datasets. Non-mob represents non-mobilizable plasmid. (B) The host range of classified plasmid genomes in the genus level. Different colors represent different genera, and genera accounting for less than 5% of the total abundance are grouped under the category ‘other’.

Additionally, almost all MOBM type plasmid genomes were present in *Clostridium* and *Enterocloster*, and some species in *Clostridium* could cause various diseases [52]. MOBQ demonstrated a broader host range, including *Acinetobacter, Agrobacterium, Escherichia, Rhizobium, Lactiplantibacillus* and *Staphylococcus*. Non-mobilizable plasmids were detected in the majority of bacteria. These results illustrated the relationship between different MOB types and their host range, and it demonstrated that MOB typing for plasmid fragments is feasible in the absence of relaxases.

### 3.2. Overall performance of MOBFinder

We used the *accuracy, kappa* and *runtime* to evaluate the overall performance of MOBFinder. Compare to MOB-suite, the *accuracy* of MOBFinder ranged from 70% to 77%, which exhibited a significant improvement of at least 59% (Figure 3A). The *kappa* of MOBFinder ranged between 67% and 75% and was approximately 65% higher than MOB-suite (Figure 3B). Moreover, MOBFinder exhibited a significantly shorter *runtime* in the test dataset, with a more gradual increase trend (Figure 3C). In general, MOBFinder demonstrated a consistent performance improvement as the sequence length increased.

**Figure 3.**
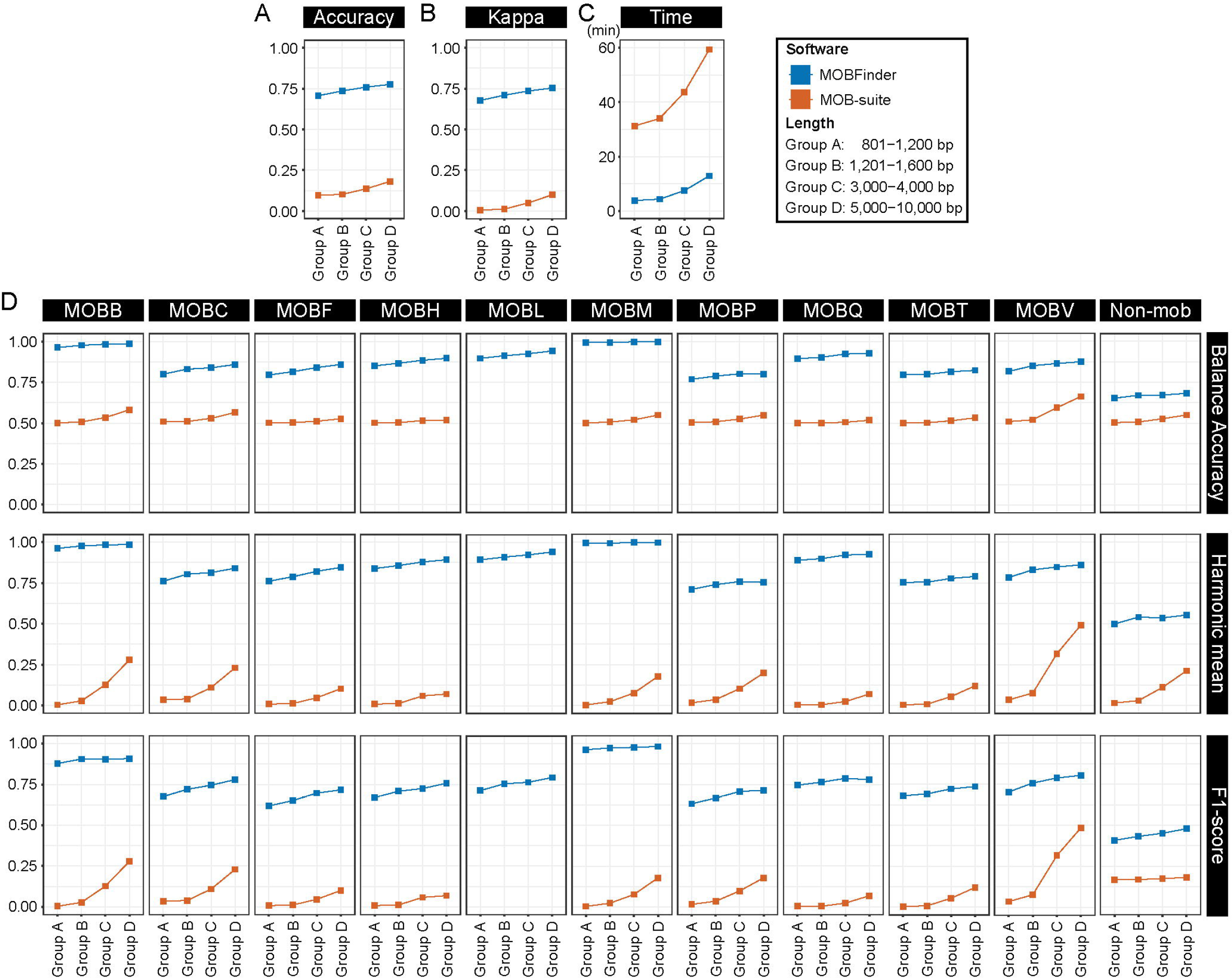
Overall performance of MOBFinder and comparison to MOB-suite. (A-C) The performance evaluation and comparison between MOBFinder and MOB-suite using *accuracy* (A), *kappa* (B) and *runtime* (C). MOBFinder is represented using blue squares and lines, and MOB-suite is represented using red squares and lines. In test datasets, four length groups were generated: Group A: 801-1200 bp, Group B: 1201-1600 bp, Group C: 3000-4000 bp and Group D: 500-10000 bp. (D) For each MOB type, the *balanced accuracy, harmonic mean* and *F1-score* were used to assess the performance of MOBFinder and compared to MOB-suite.

### 3.3. Evaluation in each MOB category

To evaluate the discrimination ability of MOBFinder in each MOB type, we calculated the *balanced accuracy, harmonic mean* and *F1-score* using the test dataset (Figure 3D). In MOBB and MOBM, MOBFinder demonstrated the highest performance, while its ability to identify non-mob class is comparatively lower than other classes. In MOBM, the *balanced accuracy* and *harmonic mean* could reach 99%, and *F1-score* exceeded 96% in all length groups. In non-mob, the *balanced accuracy* was 65%, the *harmonic mean* was 49% and the *F1-score* was 40%. Compared to MOB-suite, MOBFinder exhibited much better performance in predicting all MOB classes. Even in non-mob, MOBFinder showed approximately 13% improvement in *balanced accuracy*, 17% in *harmonic mean* and 24% in *F1-score*.

The receiver operating characteristic (ROC) curve exhibited the performance of MOBFinder in all MOB categories, and the area under the curve (AUC) was calculated to quantify the performance (Figure 4). We found all AUCs were greater than 0.8, which indicated MOBFinder could effectively distinguish positive and negative samples in each MOB class. The AUC values were higher than 0.9 in most MOB types, while in MOBT and non-mob were less than 0.9. The performance differences in identifying each MOB type might be attributed to the varying host ranges and sequence features. Additionally, the imbalance in the training dataset for each MOB type may also be a primary factor contributing to the performance disparities.

**Figure 4.**
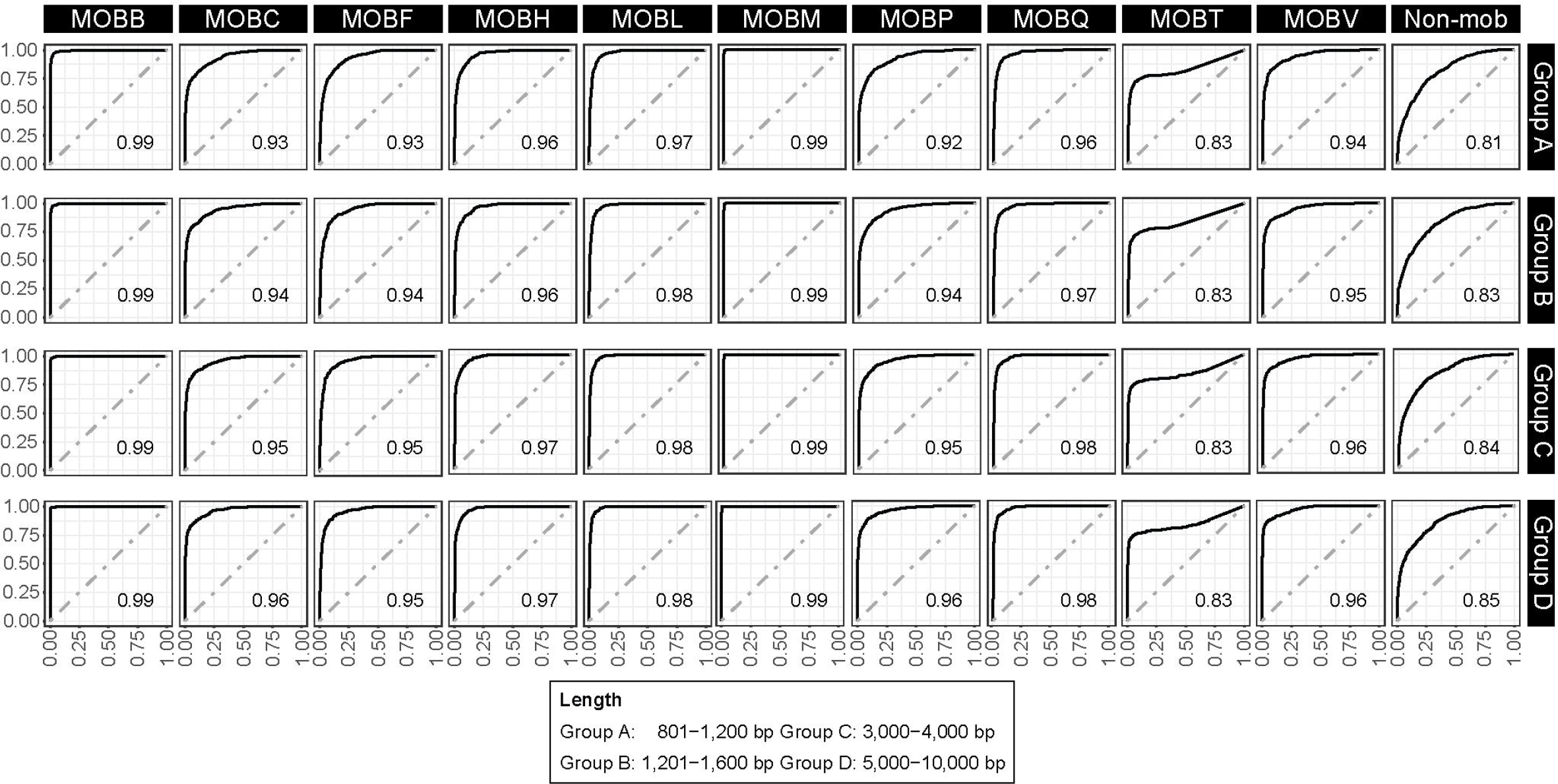
The ROC curves and AUC values of MOBFinder. The ROC curves were plotted using the output scores of MOBFinder, and the AUC values were calculated to quantify the performance of MOBFinder in each MOB class.

### 3.4. Application in T2D metagenomic analysis

Upon analysis of fecal samples in metagenomic studies, antibiotic resistance pathways were found to be enriched in patients with T2D [40]. The precise mechanism, however, remained elusive. We applied MOBFinder to analyze real T2D metagenomic data [39]. After preprocessing and assembly, 2,217,064 metagenomic fragments were generated, and plasmid assemblies were identified using PPR-Meta. Subsequently, the plasmid fragments were clustered into 55 bins and annotated using MOBFinder. By employing MOBFinder, we assigned 2 bins to the MOBF class, 8 bins to MOBL, 17 bins to MOBQ, and identified 28 bins as non-mob (Figure 5A). Furthermore, we detected 19 bins that exhibited significant difference between the T2D group and the control group. Among them, 1 bin was classified as MOBF, 2 bins as MOBL, 9 bins as MOBQ, and 7 bins as non-mob. Furthermore, MOBF is widely present in *Escherichia* and *Klebsiella* (Figure 2B), and some strains in *Klebsiella* is resistant to multiple antibiotics, including carbapenems [53]. Consequently, MOBF-type plasmids might contribute to the antibiotic resistance enrichment in T2D.

**Figure 5.**
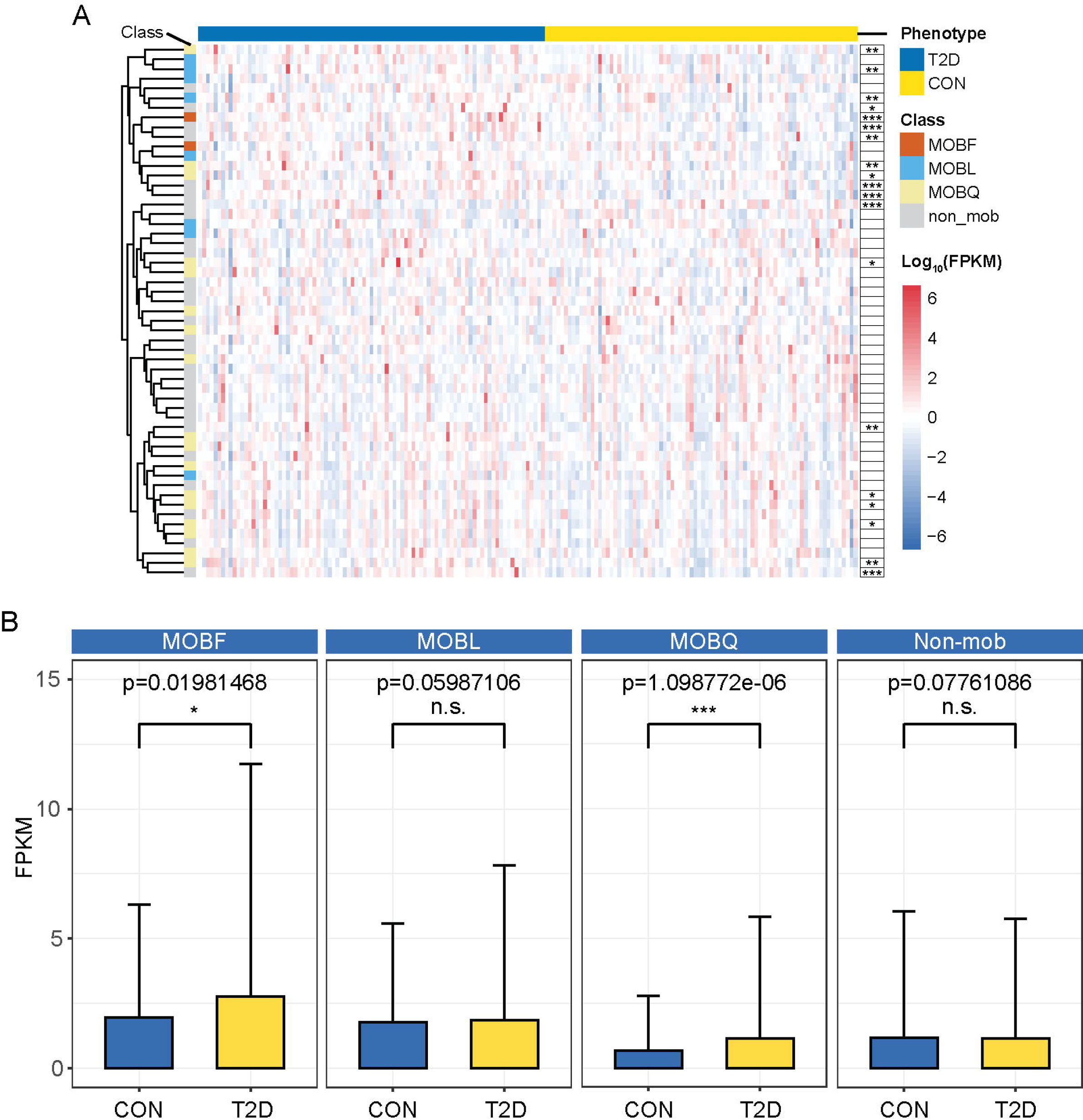
The annotation of T2D-related plasmid bins using MOBFinder. (A) Heatmap of plasmid bins between T2D and control. Each column represents a sample, and each row represents a plasmid bin. For each column, the color blue represents the T2D group, and the color yellow represents the control group. The four distinct colors on the y-axis represent identified four different MOB types using MOBFinder. (B) The abundance comparison the four identified MOB types between T2D and control. The *p-value* was calculated using Wilcoxon signed-rank two-sided test. (“*” represents *p-value* < 0.01, “**” represents *p-value* < 0.05 and “***” represents *p-value* < 0.001.)

To figure out whether MOBF-type plasmids was associated with the enrichment of antibiotic resistance in T2D, we compared the average abundance between the T2D group and the control group for each MOB type (Figure 5B). We indeed observed a significant increase in the abundance of MOBF and MOBQ within the T2D group, and these two types of MOB plasmids can be transferred among multiple bacterial species. This suggested that the increase of MOBF-type and MOBQ-type plasmids could potentially raise the risk of infection among individuals with T2D. Subsequently, we used four databases [46-49] to detect drug resistance genes in four MOB types (Figure 6). Among them, the number of identified drug resistance genes in the MOBF-type plasmids was significantly higher than the other three MOB types. This indicates that MOBF-type plasmids could carry much more drug resistance genes compared to other MOB classes. These findings suggested that the increase of MOBF-type plasmids could result in more bacteria acquiring drug resistance genes, thereby leading to the enrichment of resistance pathways in T2D. In summary, our results demonstrated the utility of MOBFinder for annotation of plasmid fragments in metagenomes, shedding light on the mechanisms fueling the emergence of antibiotic resistance enrichment in metagenomic analysis.

**Figure 6.**
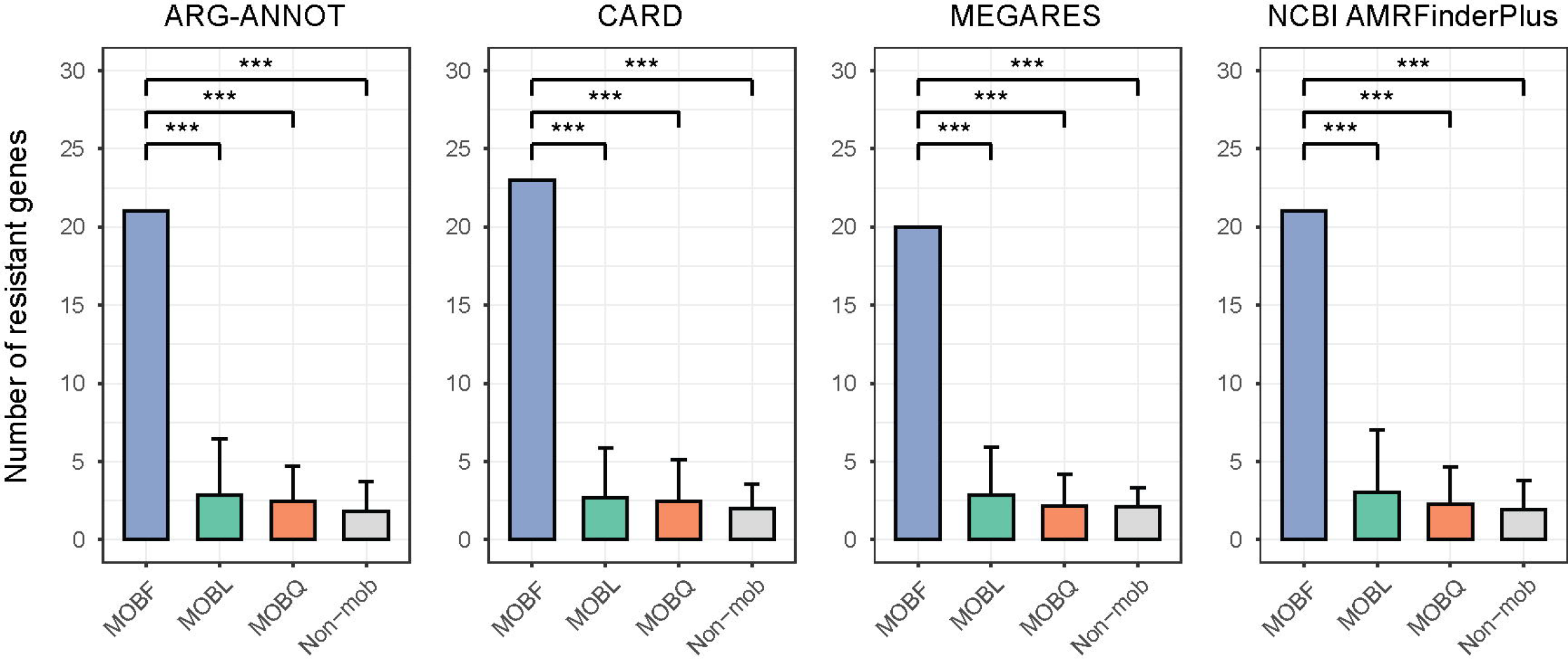
Comparison of resistant genes among different MOB types. Four databases were used to identify resistant gene within each MOB type, and the *p-value* was calculated using Tukey’s Honest Significant Difference test. (“*” represents *p-value* < 0.01, “**” represents *p-value* < 0.05 and “***” represents *p-value* < 0.001.)

### 3.5. The usage of MOBFinder

MOBFinder can predict the MOB type of plasmid fragments and bins in metagenomics. For plasmid metagenomic fragments, MOBFinder takes a FASTA file as input. The output file consists of 13 columns. The first column represents the fragment ID, the second column displays the predicted MOB type and columns three to thirteen represent the scores for each MOB class, namely MOBB, MOBC, MOBF, MOBH, MOBL, MOBM, MOBP, MOBQ, MOBT, MOBV, and non-mob.

For plasmid metagenomic bins, MOBFinder requires two input files: a FASTA file containing the plasmid fragments and a meta table that records the mapping between plasmid fragment IDs and bin IDs. The output results are similar to the output of plasmid fragments. The first column is the plasmid bin’s id. The second column is the predicted MOB class of plasmid bins. The other columns were MOB scores of different MOB categories produced by MOBFinder.

## 4. Discussion

In this paper, we developed MOBFidner based on a language model [30, 31] and the random forest. Using the relaxase-alignment method, plasmid genomes were classified into distinct MOB categories. Our analysis revealed substantial variations in parameters such as the number, average length, and GC content across various MOB types. Additionally, there are noteworthy differences in the host range among different MOB classes. These results suggested the potential of utilizing sequence features from different MOB types for plasmid metagenomic fragment MOB typing. To characterize the plasmids within each MOB type, we used skip-gram to generate word vectors. MOBFinder, integrating multiple random forest models, demonstrated superior overall performance compared to existing tools. Specifically, for each MOB category, MOBFinder exhibited significant improvements in *balanced accuracy, harmonic mean* and *F1-score*, with values reaching up to 99% in the MOBM category.

Although MOBFinder demonstrated commendable performance, its efficacy in some MOB types was lower than in others, such as in MOBP, MOBT and non-mob. The host range of these MOB categories is more diverse and complex compared to others (Figure 2B), potentially encompasses additional features that we have not captured. Furthermore, there is a possibility that some MOB types remain undiscovered, thus necessitating additional experimental validation.

Interestingly, through our analysis of T2D metagenomic sequencing data [39], we noted a significant increase in MOBF and MOBQ type plasmids within T2D patients. Moreover, we found that the drug resistance genes were enriched in the MOBF class, which were also present in *Klebsiella* that is associated with the spread of multidrug resistance. Although previous analysis of metagenomic data from patients with T2D found the drug resistance pathways enriched [40], our analysis revealed a potential reason behind this phenomenon: the increased abundance of MOBF-type plasmids in the gut of individuals with T2D may lead to the dissemination of more antibiotic resistance genes, resulting in the enrichment of the antibiotic resistance pathway.

At present, a large amount of human metagenomic data derived from second-generation sequencing populates various databases. However, our understanding of the functions of numerous disease-linked microbial sequences remains limited, attributed to the incomplete nature of metagenomic fragments. The development of MOBFinder enables the MOB annotation for plasmid fragments in metagenomics, and provides a powerful tool to investigate the transmission mechanisms of plasmid-mediated antibiotic resistance genes and virulence factors.

## 5. Conclusions

In summary, we have developed MOBFinder as a tool for MOB typing of plasmid fragments and bins in metagenomic data. Our analysis of classified plasmid genomes unveiled notable differences in sequence characteristics and host range across various MOB types. Based on this, we employed a language model to extract the sequence features specific to each MOB type and represented them using word vectors. Additionally, we boosted prediction accuracy by training and integrating several random forest classification models. MOBFinder surpassed the existing tool in the performance tests and was successful in detecting an increase in MOBF type plasmids in T2D patients from the T2D metagenomic data analysis. Importantly, the MOBF type plasmids harbor potential drug-resistant genes, thus offering an explanation for the observed antibiotic resistance in T2D individuals. This suggests MOBFinder’s potential in aiding the formulation of specific medications to curb drug resistance transmission. We anticipate that MOBFinder will be a powerful tool for the analysis of plasmid-medicated transmission.

## Availability of Source Code and Requirements

- Project name: MOBFinder
- Project homepage: https://github.com/FengTaoSMU/MOBFinder
- Operating system(s): Linux
- Programming language: Python, R script
- Other requirements: BLAST, biopython
- License: GPL-3.0
- RRID: SCR_024451

## Availability of Supporting Data

Snapshots of our code and other data further supporting this work are openly available in the GigaScience repository, GigaDB

## Abbreviations

MOB: mobilization; Rep: replicon; Inc: incompatibility; MPF: mate-pair formation; non-mob: non-mobilizable; T2D: type 2 diabetes; FPKM: fragments per kilobase per million; TPR: true positive rate; TNR: true negative rate; Sn: sensitivity; Sp: specificity; ROC: the receiver operating characteristic; AUC: the area under the curve; SRA: short read archive; NCBI: National Center for Biotechnology Information; PlasTax-PCR: PLASmid TAXonomic PCR; PBRT: PCR-Based Replicon Typing; DPMT: Degenerate Primer MOB Typing.

## Authors’ Contributions

TF, ZCF and HWZ proposed and designed this work. TF and ZCF developed and optimized the software. TF, ZCF, SFW and HWZ wrote and revised the manuscript.

## Supplementary data

### Supplementary Material

**Supplementary Table 1**. The accessions list of classified plasmid genomes.

**Supplementary Table 2**. The list of metagenomic samples used in our analysis.

**Supplementary Figure 1**. MOB typing using MOB-suite. Single-class represents plasmid genomes classified into one MOB type, multi-class represents plasmid genomes classified into more than one MOB categories and non-mob represents non-mobilizable plasmids.

